## Supplementary figures and images for "Integration of whole genome sequencing and transcriptomics reveals a complex picture of insecticide resistance in the major malaria vector *Anopheles coluzzii*"

### Supplementary Figure 1

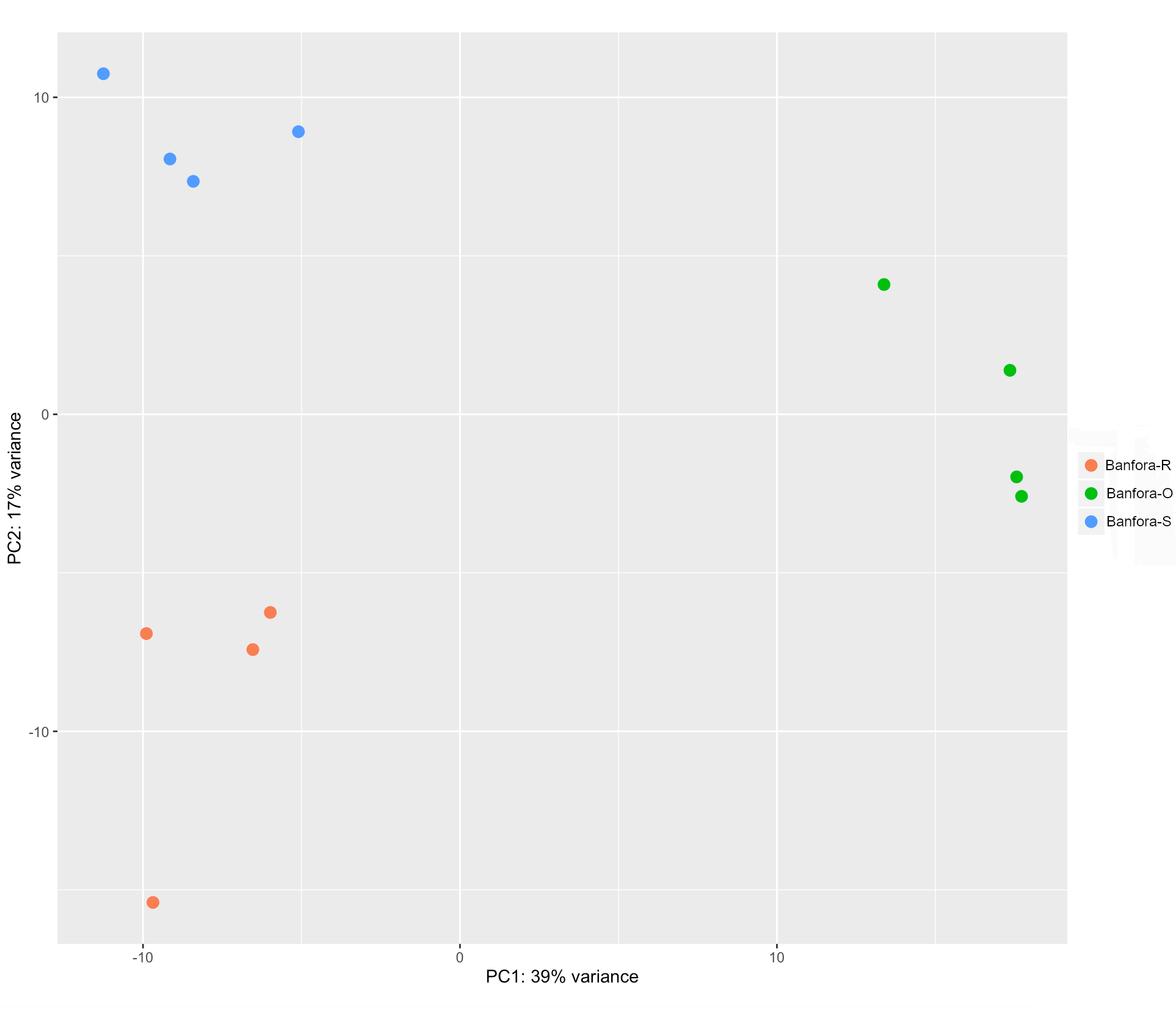

### Supplementary Figure 2

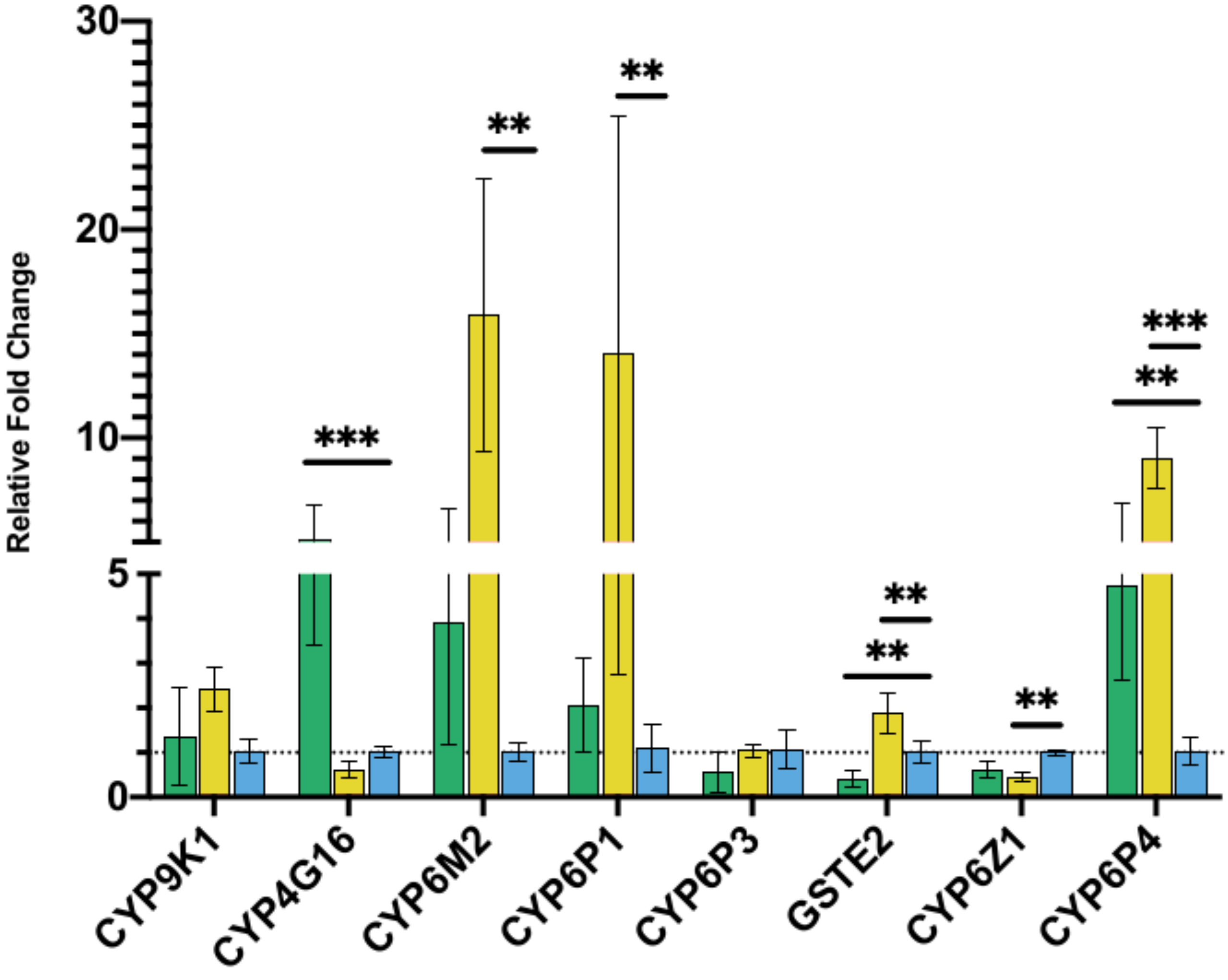

### Supplementary Figure 3

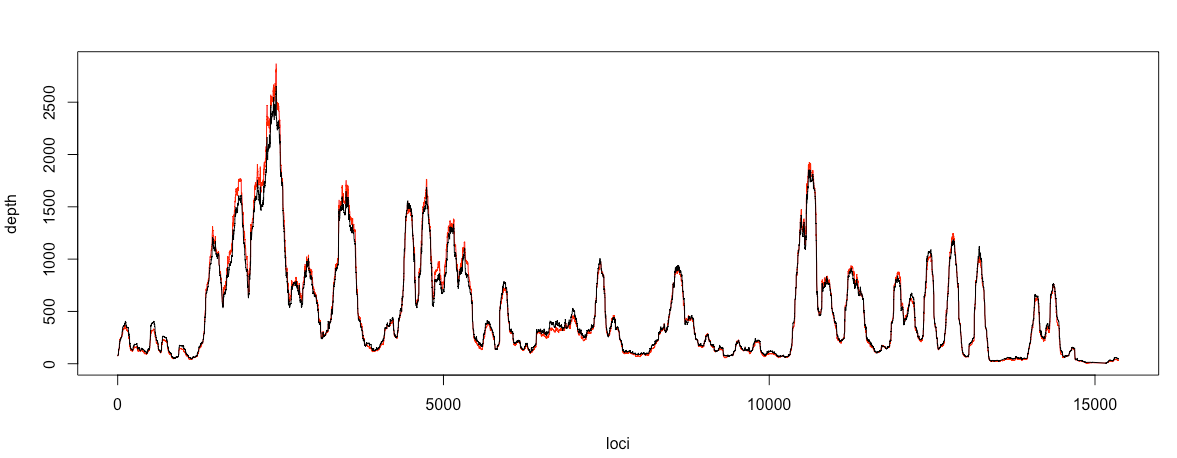

### Supplementary Figure 4

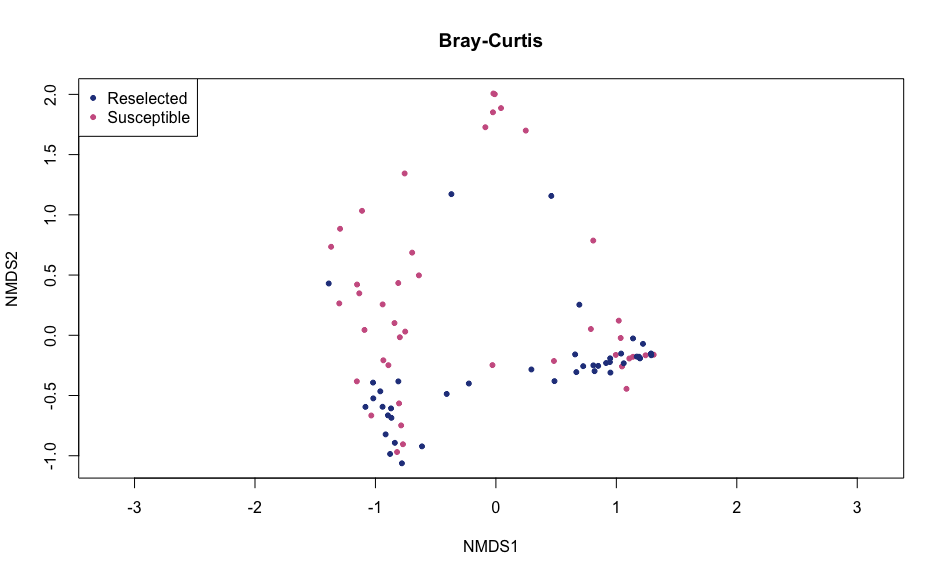

### Supplementary Figure 5

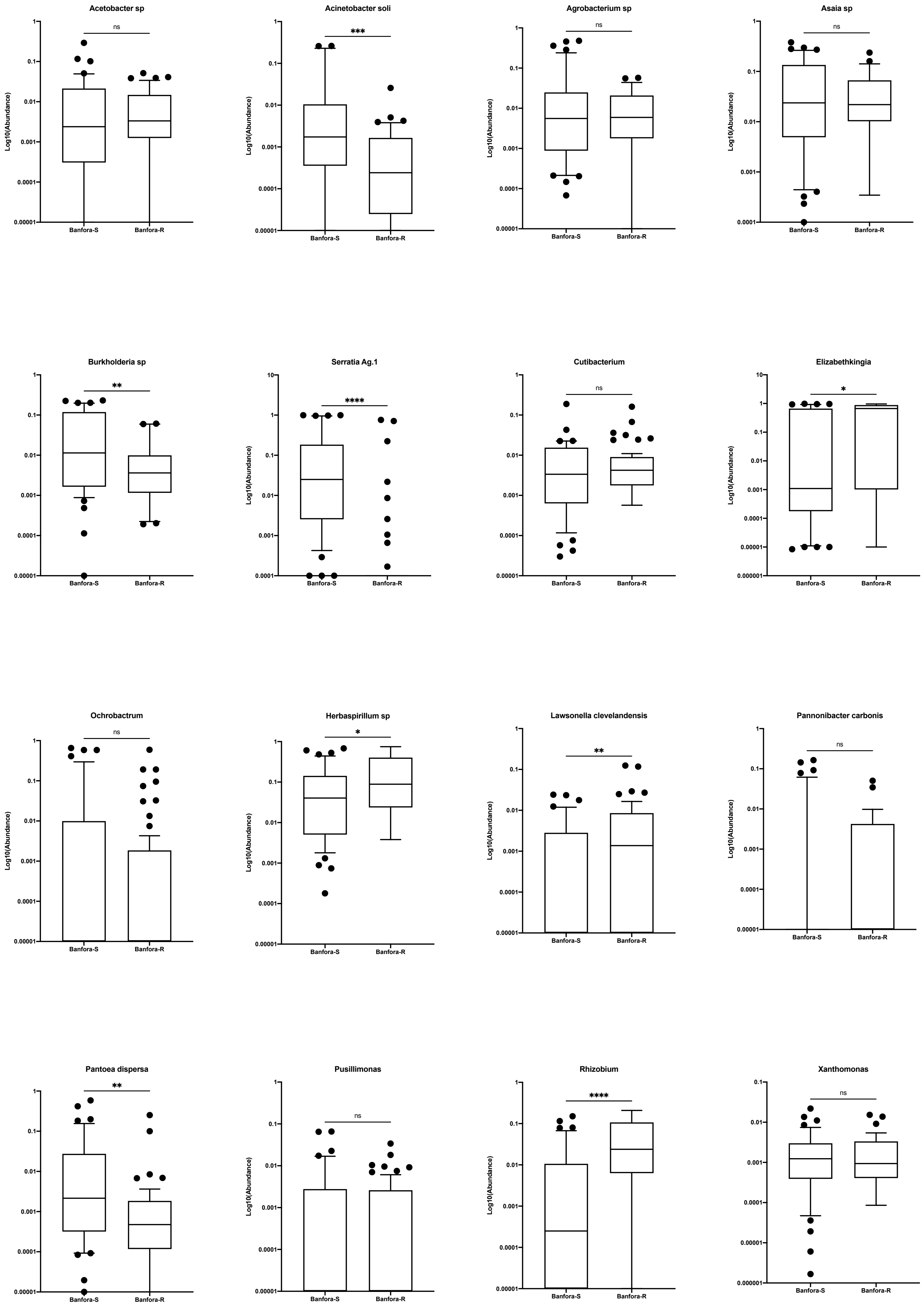

### Supplementary Figure 6

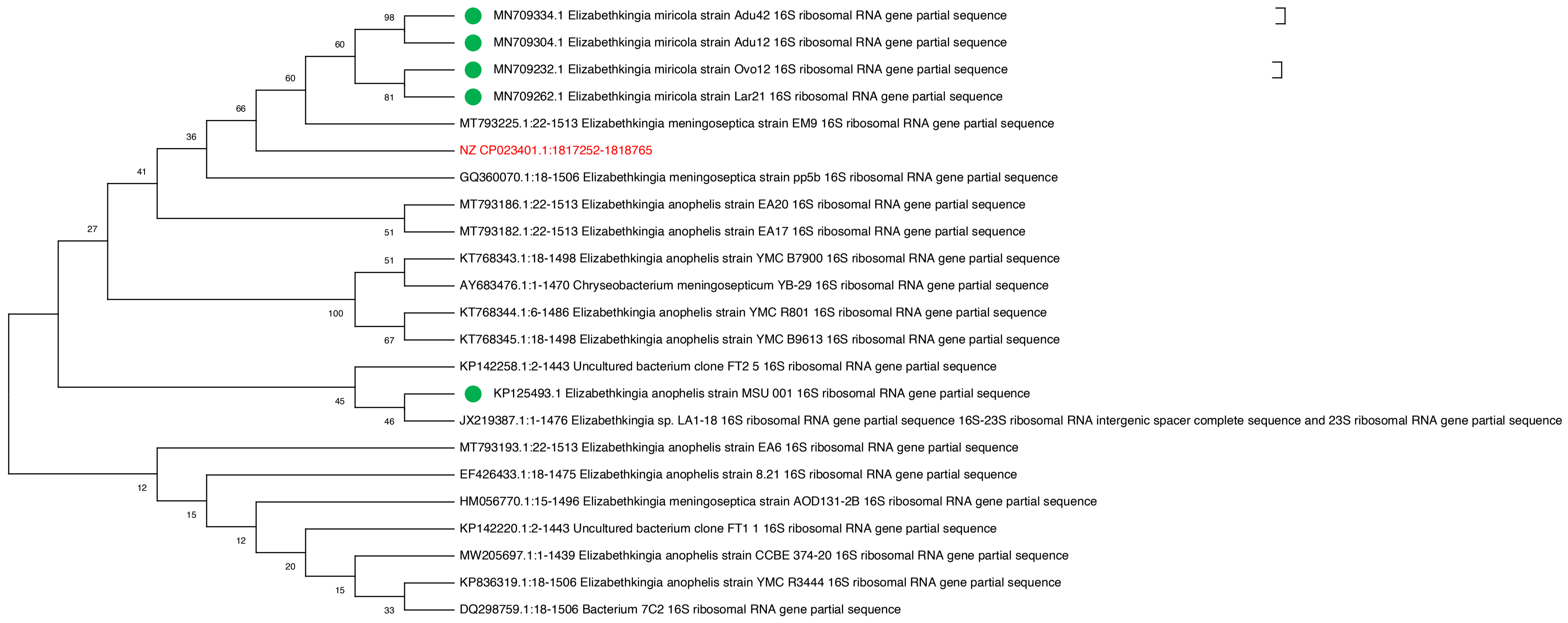

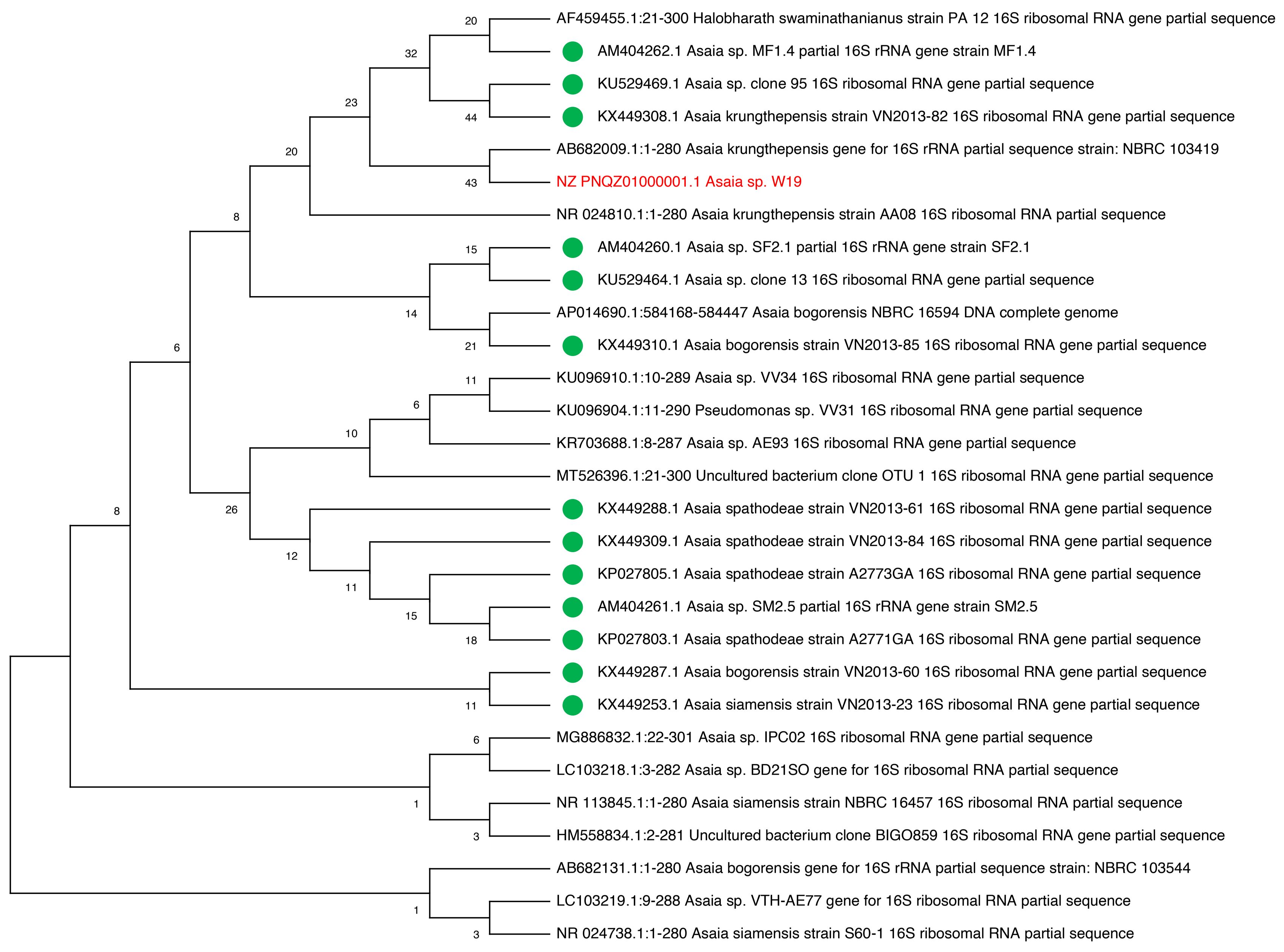

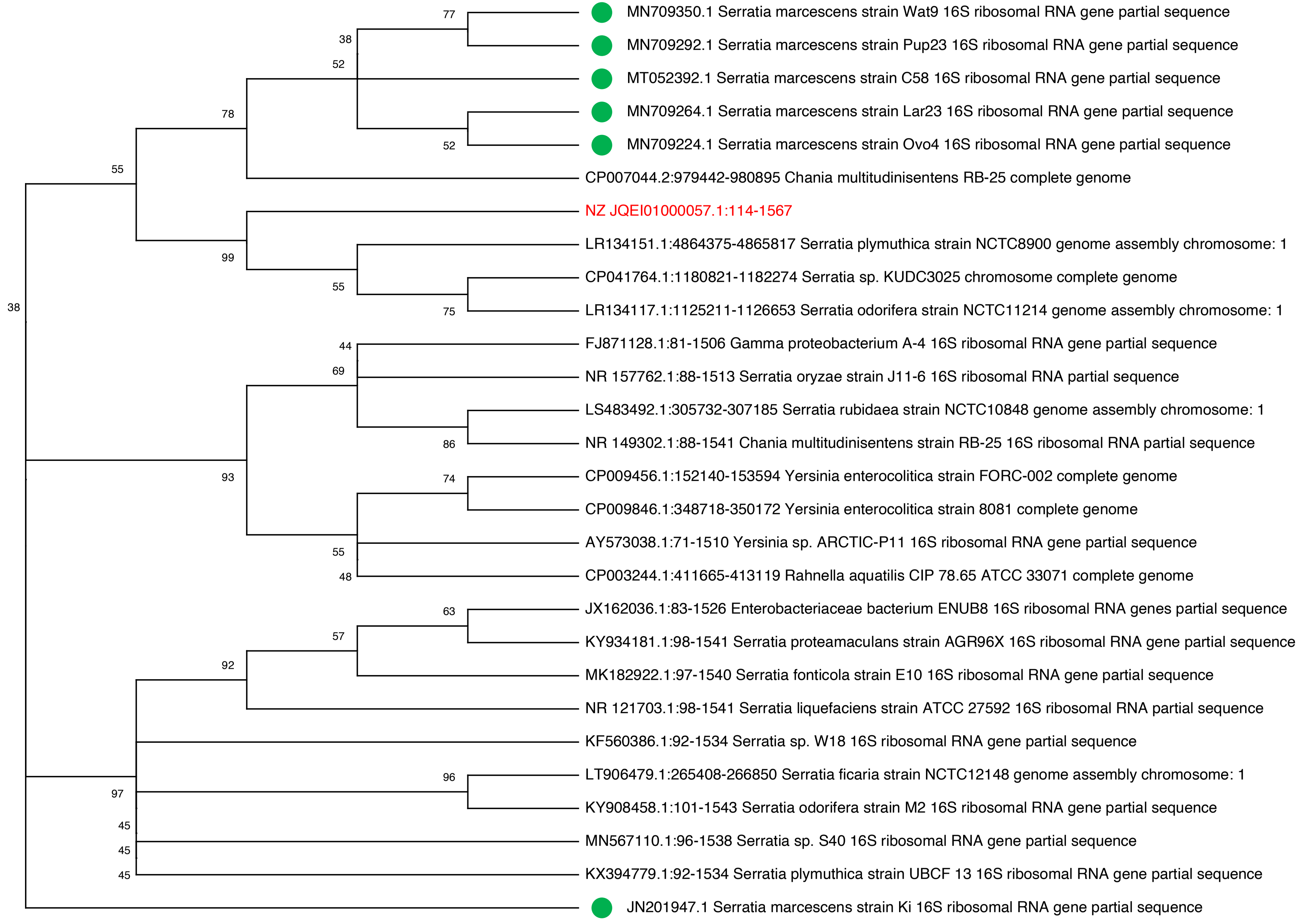
