## Appendix 1 for "Integration of whole genome sequencing and transcriptomics reveals a complex picture of insecticide resistance in the major malaria vector *Anopheles coluzzii*"

Chromosome 2R

The first peak on 2R (a) corresponds to a 3.4Mbp block and includes 188 genes, encompassing the entire CYP325 cluster, two of which are significantly down-regulated, CYP325D2 and CYP325D1. Three missense mutations in the immune gene LRIM5 (AGAP027997), and one in each of AGAP002345, AGAP002352 and AGAP002354 are within this region. Of the 188 genes, 69 are differentially expressed, with 35 up-regulated in ? and *Gnmt,* previously linked with the *Maf-S* pathway showing the highest fold change (3x).

Chromosome 2L

A small 39kb region (c) at the start of 2L contains just one gene, of unknown function (AGAP028432) that is significantly down regulated (p = 6.25e-11; FC = 0.43) and has a GO term related to protein binding. Within this 2Ld region, there are 331 genes of which 141 are differentially expressed. Amongst these were 86 up-regulated genes, including three up-regulated prophenoloxidases and two up-regulated transcription factors, IMD and mbf1, recently linked to insecticide resistance ^34^ but interestingly *kdr* and three potassium gated channels AGAP00417 to AGAP00419 were down-regulated. Two further small peaks are present on 2L including a 222kb region (e) and an 822kb region (f); the former contains seven genes whilst the latter has 68.

Chromosome 3R

Two blocks of peaks are present on 3R, the first consists of 1.04Mb encompassing 82 genes and containing two missense variants in AGAP007880 a 3-ketoacyl-CoA reductase involved in fatty acid elongation and AGAP007878, a gene of unknown function. The second peak on 3R

corresponds to a 1.3Mbp block containing the ABCC, CPLCG and CPLCW clusters. Within these clusters ABCC8 and ABCC11 are significantly upregulated (2.37x and 3.01x), of the CPCL family, only CPCLG5 is differentially expressed, the gene in this family previously linked with pyrethroid resistance (3.84x).

X Chromosome

The first peak on the X chromosome block (i) covers 432kbp and includes 11 genes, with just three showing differential expression. Missense variants are present in five of the genes, one in each of AGAP000150-AGAP000153, AGAP000155 and three missense variants are present in AGAP013500. AGAP000155, a DNA repair related protein is significantly overexpressed in the RNAseq. The second peak on the X chromosome is 105kb, with all SNPs occurring in intragenic regions, impacting one gene AGAP001094 which is highly significantly down regulated (p = 5.72e-10, FC = 0.43). AGAP001094 is the homolog of the Drosophila transcription factor *runt,* involved in nervous development and maintenance. Of the known *runt* interactors in *Drosophila*, 60% are differentially expressed in the Banfora-R line.
